# Global Proteomics Investigation of SAMT-247 Targets: An Antiviral Thioester that Acetylates Zinc Finger Proteins

**DOI:** 10.64898/2026.04.28.721345

**Authors:** Connor P. Jewell, Andrew J. Perciaccante, Kate Brown, Tapan K. Maity, Jerry C. Dinan, Massimiliano Bissa, Mohammad Arif Rahman, Genoveffa Franchini, Daniel H. Appella, Lisa M. Jenkins

**Author notes:** These authors contributed equally (Connor P. Jewell, Andrew J. Perciaccante). Corresponding author: Lisa M. Jenkins, Laboratory of Cell Biology, National Cancer Institute, NIH, 37 Convent Drive, Room 2068, Bethesda, MD 20892.

## Abstract

Covalent modification of target proteins is a well-established mechanism of action for small molecule inhibitors. Cysteine residues in particular have been exploited for their reactivity toward electrophilic molecules. SAMT-247 is a mercaptobenzamide thioester that covalently acetylates cysteines in the zinc-coordinating domains of the HIV nucleocapsid protein. This SAMT-247-promoted reaction leads to loss of zinc binding by the protein, with concomitant loss of protein structure and function. Although it has low cytotoxicity in animal models, recent studies have indicated that it affects other protein targets in uninfected cells, for example leading to increased immune cell functions. In this study, global proteomics approaches have been used to better understand other protein targets of SAMT-247. Minimal effects are observed when unstimulated THP-1 monocyte cells were treated with SAMT-247. In contrast, thermal proteome profiling identified 170 proteins with altered thermal stability when THP-1 cells were stimulated with phorbol 12-myristate 13-acetate/Ionomycin (PMA/Iono) before SAMT-247 treatment. Among the affected proteins, 81 contain a zinc-coordinating domain and/or have been shown to have a reactive cysteine residue. Among these, several play a role in cellular metabolism, and Seahorse assays demonstrated that SAMT-247 significantly increased the anti-metabolic and pro-glycolytic effect of PMA/Iono in THP-1 cells. Two of the most-affected proteins were ZC3H7A, a microRNA-binding protein with four zinc finger domains, and MGMT, a DNA damage repair protein with a reactive cysteine. Both proteins were modified by SAMT-247 when tested alone or in the presence of THP-1 cell lysate, indicating that they are bona fide targets of the inhibitor. The low activity of SAMT-247 in unstimulated THP-1 cells is consistent with its low cytotoxicity. The increased effects of SAMT-247 in stimulated immune cells suggests that this molecule could be developed to target diseases other than HIV.

## Introduction

Covalent modification of protein targets is an important mechanism of action for many small molecule drugs. Because covalent modification of a protein is permanent and irreversible, a molecule with weak binding affinity can still achieve good therapeutic efficacy using this mechanism. Furthermore, covalent inhibitors may avoid problems with resistance and promote the usage of lower dosages.^1^ Several well-known drugs function via covalent inhibition, including penicillin and aspirin.^2^ Covalent modification has been shown to facilitate targeting of “undruggable” proteins, such as KRAS,^3^ and to allow non-specific targeting of structures such as heterochromatin to prevent mitotic division, a new approach to the treatment of cancer.^4^ However, these inhibitors can have detrimental off-target effects that result in unexpected toxicity or hypersensitivity. Thus, it is critical to understand the selectivity of any covalent inhibitor.

SAMT-247 is a covalent inhibitor designed to target the nucleocapsid (NC) protein of the HIV. SAMT-247 preferentially reacts with Cys36 of the C-terminal zinc-binding domain of NC, covalently transferring an acetyl group to the cysteine sidechain and releasing a free thiol (Figure 1).^5, 6^ This intermolecular transfer is followed by intramolecular transfer of the acetyl group from the cysteine thiol to a proximal lysine sidechain, leading to irreversible chemical acetylation of the primary amine in the sidechain. Both the cysteine and lysine modifications disrupt zinc coordination and result in loss of protein structure. Once the initial reactions occur, multiple sites within the both zinc-binding domains can be modified due to overall loss of the structure and exposure of cysteine residues.^6, 7^ Intriguingly, the free thiol that is released following reaction with NC is acylated intracellularly to re-form an active thioester, therefore allowing the inhibitor to be intracellularly regenerated.^6^ Thus, rather than a single 1:1 reaction model, SAMT-247 can potentially inhibit many molecules of its target.

**Figure 1.**
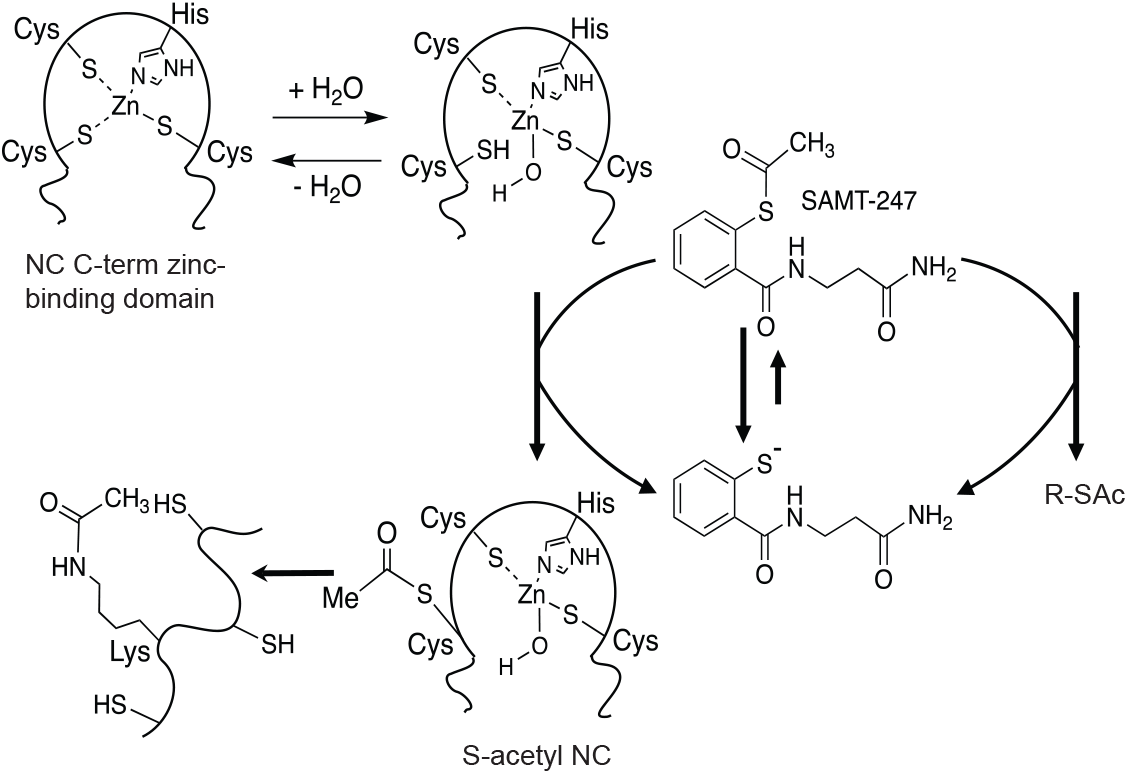
Schematic of SAMT-247 reaction mechanism against HIV-1 NC protein. Initial modification of cysteine residues in the zinc-finger domain by SAMT-247 results in transfer of an acetyl group from SAMT-247 to the thiol (SH) side chain. This is followed by intramolecular transfer of the acetyl group to nearby lysine residues forming a stable modification of the protein.

SAMT-247 is an effective inhibitor of HIV in multiple cellular and animal models.^8-11^ Most recently, SAMT-247 has been shown to combine with an SIV vaccine in macaques to confer increased protection against infection as compared with the vaccine alone.^9^ Concomitantly, SAMT-247 has low cellular toxicity in all systems in which it has been studied. While this would seemingly suggest high overall specificity of reaction, *in vitro* studies with analogs of SAMT-247 have demonstrated that multiple different classes of zinc-coordinating domains can be targeted similarly to HIV NC, indicating that SAMT-247 activity may be more broad.^12^ Consistent with this, *in vitro* and *ex vivo* studies demonstrated that SAMT-247 has immunomodulatory effects on diverse systemic and mucosal immune cells.^9^

To better understand potential targets of SAMT-247, global proteomics approaches have been used. These provide an unbiased method to look for effects of SAMT-247 that could be beneficial or detrimental. Here, we have focused on the effect of SAMT-247 in immune cells, comparing unstimulated and stimulated conditions. We show that there are multiple new potential targets of SAMT-247 and reaction of two specific proteins was demonstrated both *in vitro* and in a cellular milieu. These studies suggest new applications for this inhibitor in immune modulation and cancer.

## Experimental Methods

### Global Proteome Analysis

THP-1 cells were treated in biological triplicate with SAMT-247 (100 µM) or DMSO (vehicle control) for 12 h before harvesting. Pellets were lysed in 50mM Tris HCl pH 7.5, 120 mM NaCl, 5mM EDTA, 0.5% NP40, supplemented with protease inhibitor. Samples were then reduced by addition of 100 mM dithiothreitol (10 mM final concentration) at 56 °C for 1 h. Samples were cooled to room temperature and alkylated by addition of 400 mM Iodoacetamide (20 mM final concentration) at room temperature for 30 minutes in the dark. Following alkylation, samples were diluted 4-fold with 50 mM triethylammonium bicarbonate (TEAB) digested with trypsin/Lys-C (1:125, Promega) at 37 °C overnight. Digested peptides were acidified with 20% trifluoroacetic acid (TFA) to a final concentration of 0.5% and purified using C18 desalting spin columns (Thermo Scientific) and lyophilized. Dried peptides were suspended in 5 mM TEAB and concentrations were measured using a colorimetric BCA peptide assay (Thermo Scientific). From each sample, 100 µg of peptide were labeled with TMT 6-plex reagent (Thermo Scientific) following manufacturer’s instructions. Following labeling, peptide samples were pooled and lyophilized, then offline fractionated by high pH reversed-phase separation (Thermo Scientific) with twelve elution buffers containing 0.1% triethylamine, and incremental concentrations (5%, 7.5%, 10%, 12%, 14%, 16%, 18%, 20%, 22%, 25%, 35%, and 60%) of acetonitrile. Fractions were lyophilized separately.

### Thermal Proteome Profiling

THP-1 cells were treated in biological duplicate with SAMT-247 (100 µM) or vehicle control for 12 h before harvesting and lysed with 50mM Tris HCl pH 7.5, 120 mM NaCl, 5mM EDTA, 0.5% NP40, supplemented with protease inhibitor. Duplicate cellular lysates of each condition were divided into 16 aliquots of 45 µL and transferred to PCR tubes. Each tube was heated to a designated temperature (37, 42.4, 45.6, 50.1, 53.9, 58.4, 61.7, 67 °C) for three minutes using the thermal cycler function of a Duet CFX Real-Time PCR System (BioRad). Samples were then left at room temperature for 3 additional minutes before placed on ice and transferred to 1.5 mL low protein binding Eppendorf tubes. Denatured, aggregated proteins were removed via high-speed centrifugation set at 4 °C for 20 minutes and 17,000 x g. The supernatant was collected, placed in new low protein binding tubes, and stored at −20 °C overnight.

Each sample was transferred to a low-bind tube and spun down at 4 °C for 20 minutes at 17,000 x g. Subsequently, 40 µL of supernatant was removed to new tubes for processing. Protein quantitation was performed by BCA (Thermo Fisher Scientific) and the amount of sample at 37 °C was normalized for the DMSO and SAMT-247 treatments. Each sample was diluted with 50 mM triethylammonium bicarbonate (TEAB) to a final volume of 100 µL. Reduction with 100 mM dithiothreitol (DTT) in 50 mM TEAB was performed by incubation at 57 °C for 45 minutes. Samples were then alkylated with 400 mM iodoacetamide in 50 mM TEAB at room temperature in the dark for 30 minutes. Trypsin (1:50, Thermo Fisher Scientific) digestion was carried out at 37 °C for 16 hours overnight. Digested peptides were lyophilized to dryness and then desalted using Desalting Spin Columns (Thermo Fisher Scientific). After another lyophilization, peptides were quantified using a colorimetric kit (Thermo Fisher Scientific) and TMT-labeled using a TMTpro-16plex kit (Thermo Fisher Scientific), following manufacturer’s instructions. After labeling, samples were pooled by treatment condition at equal volumes and lyophilized. Samples were then reconstituted in 0.1% trifluoroacetic acid (TFA) and fractionated using a high pH Reversed-Phase Peptide Fractionation Kit (Thermo Fisher Scientific). The provided protocol was altered for a final production of 12 gradient condition fragments at an acetonitrile percentage of 5, 7.5, 10, 12, 14, 16, 18, 20, 22, 25, 35, and 60%. Resultant fractions were separately lyophilized.

### Mass Spectrometry Analysis

For the global and TPP mass spectrometry analyses, dried peptides were resuspended in 5% acetonitrile, 0.05% TFA in water for analysis on an Orbitrap Exploris 480 (Thermo) mass spectrometer. The peptides were separated on a 75 µm × 15 cm, 3 µm Acclaim PepMap reverse phase column (Thermo) at 300 nL/min using an UltiMate 3000 RSLCnano HPLC (Thermo) and eluted directly into the mass spectrometer. Parent full-scan mass spectra collected in the Orbitrap mass analyzer set to acquire data at 120,000 FWHM resolution; fragments were isolated with a 0.7 m/z isolation window, fragmented by HCD at 30%, and product spectra collected at 45,000 FWHM resolution. For the TPP analyses, duplicate or triplicate technical replicate injections were performed in addition to the biological replicates.

Global proteome data were searched using Proteome Discoverer 3.0 (Thermo Scientific) against the human database from Uniprot using SequestHT. The search was limited to tryptic peptides, with maximally two missed cleavages allowed. Cysteine carbamidomethylation and TMT modification of peptide N-termini and lysines were set as a fixed modifications; methionine oxidation was set as a variable modification. The precursor mass tolerance was 10 ppm, and the fragment mass tolerance was 0.02 Da. The Percolator node was used to score and rank peptide matches using a 1% false discovery rate. TMT quantitation was performed using the Reporter Ions Quantifier nodes with correction of the values for lot-specific TMT reagent isotopic impurities.

TPP data were searched using Proteome Discoverer 3.0 (Thermo Scientific) against the human database from Uniprot using SequestHT. The search was limited to tryptic peptides, with maximally two missed cleavages allowed. Cysteine carbamidomethylation and TMTpro modification of peptide N-termini and lysines were set as a fixed modifications; methionine oxidation was set as a variable modification. The precursor mass tolerance was 10 ppm, and the fragment mass tolerance was 0.02 Da. The Percolator node was used to score and rank peptide matches using a 1% false discovery rate. TMT quantitation was performed using the Reporter Ions Quantifier nodes. Protein abundance data were analyzed using a custom R-based pipeline incorporating components of MineCETSA version 1.1.1^13^ and the nonparametric analysis of response curves (NPARC) framework version 1.12.0^14^. All analyses were developed and executed in RStudio using R version 4.3.1. Soluble protein abundance values were scaled using MineCETSA and used to construct thermal denaturation profiles across the temperature gradient. For each protein, nonparametric sigmoidal response curves were fitted under null (shared curve across conditions) and alternative (condition-specific curves) hypotheses. Statistical significance of treatment-dependent thermal stability changes was assessed via calculation of an F-statistic based on differences in model residuals. Resulting p-values were adjusted for multiple hypothesis testing using the Benjamini–Hochberg procedure, with a false discovery rate threshold of 5%. Proteins only present in one replicate were not considered in further analysis.

### Structural and interactome analyses

Protein domain annotation was performed using UniProt. Protein–protein interaction networks were generated using STRING database version 12.0,^15^ with the full STRING network selected and a minimum required interaction score set to medium confidence (0.400). Druggability assessment was conducted using CysDB version 1.5^16^ and associated datasets, and proteins annotated as ligandable were included in downstream analyses. When comparing melting curves, Z-score similarities were calculated using parameters derived from significantly shifted proteins only. We highlight results with z-score differences of the residual sum of squares (RSS) in the null and alternative hypothesis of less than 0.1.

### Extracellular Flux Analysis

THP-1 cells were maintained in RPMI supplemented with 10% FBS (HyClone), 1% Penicillin– Streptomycin (Sigma) and 0.1% Amphotericin B (Gibco). Treatment with SAMT-247, PMA/Iono or a combination were carried out in complete RPMI media. SAMT-247 treatment was performed using 100 nM compound for 24 h, PMA/Iono treatment was performed using 1X compound for 18 h. DMSO was used as a carrier control for all treatments.

Cellular respiration was measured using the Seahorse XF96 Flux Analyzer and the XF Cell Mito Stress Test kit according to the manufacturer’s instructions (Agilent). Cells were seeded onto Poly-D-Lysine-coated plates. The oxygen consumption rate (OCR) was measured under basal conditions and in response to the ATP synthetase inhibitor oligomycin (2.5 mM), the electron transport chain (ETC) accelerator, FCCP (1 mM), and finally the ETC complex 1 and 3 inhibitors antimycin A and rotenone (2.5 mM each). Addition of these compounds allow for the following calculations: basal respiration was calculated from the average OCR data over the initial 25 minutes after subtracting the average non-mitochondrial respiration determined after addition of Antimycin A and rotenone; ATP production was calculated by the change in OCR before and after injection of oligomycin; maximal respiratory capacity was calculated after injection of FCCP as the maximum OCR reading minus the non-mitochondrial respiration rate; spare capacity was calculated by subtracting the basal respiration from the maximal respiration rate; proton leak was calculated by subtracting the non-mitochondrial respiration from the ATP production.

Cellular glycolytic function was measured using the Seahorse XF96 Flux Analyzer and the XF Glucose Stress Test kit according to the manufacturer’s instructions (Agilent). Cells were seeded onto Poly-D-Lysine-coated plates. The Extracellular Acidification Rate (ECAR) was measured under basal conditions and in response to glucose (10 mM), the ATP synthase inhibitor oligomycin (2.5 mM), and finally the glycolysis inhibitor 2-deoxy-D-glucose (2DG, 100 mM). Addition of these compounds allows the following parameters to be calculated: basal ECAR was calculated from the average ECAR data over the initial 25 minutes by subtracting the average non-glycolytic acidification after adding 2DG; glycolysis was estimated as the increase in basal ECAR after the addition of glucose; glycolytic capacity was calculated as the maximum ECAR reading after the addition of oligomycin subtracting non-glycolytic acidification; glycolytic reserve was calculated by subtracting glycolysis from the glycolytic capacity.

Data were normalized to cell number following completion of the assay and analyzed using the XF software (Seahorse Bioscience, Agilent Technologies LDA UK Ltd). Assays were completed at least three times with five biological replicates in each experiment.

### In vitro reaction of SAMT-247 with ZC3H7A

ZC3H7A peptides representing the four zinc-coordination domains were obtained from LifeTein (New Jersey). Prior to use, peptides were refolded with zinc by titration of the solvent pH as previously described.^17^ For reaction, peptides were incubated with or without SAMT-247 at a molar ratio of 1:5 at 37 °C for 1 h. Following reaction, samples were mixed with 2% Trifluoroacetic acid (TFA), 20% acetonitrile at 3:1 ratio and purified using C18 columns (Thermo Scientific) following the manufacturer’s protocol. Briefly, C18 columns were activated with 50% methanol followed by equilibration with 0.5% TFA and 5% acetonitrile. Modified peptides were loaded onto the columns and columns were washed with the above binding buffer to remove unreacted SAMT-247. Bound peptides were then eluted with 0.5% TFA, 50% acetonitrile and lyophilized. Desalted peptides were then analyzed by mass spectrometry on an X500B Q-TOF (SCIEX) coupled to an Exion liquid chromatography system.

Recombinant ZC3H7A440-971 was purified as a His6-MBP-tevH-ZC3H7A fusion protein from baculovirus-infected insect cells and purified using immobilized metal affinity chromatography. For analysis of reaction, 10 µM ZC3H7A_440-971_ was incubated with 50 µM SAMT-247 alone or in the presence of THP-1 lysate at 37 °C for 1 hour. Control reactions were performed using DMSO as vehicle control. Following reaction in the lysate, the ZC3H7A_440-971_ was enriched from the lysate by pulldown with cobalt affinity beads (Thermo Fisher Scientific). Proteins were alkylated with iodoacetamide without prior reduction to preserve modifications on cysteine and digested with trypsin (Thermo Scientific) at an enzyme to peptide ratio of 1:50 at 37 °C overnight followed by mass spectrometry analysis on an Orbitrap Fusion (Thermo) or Orbitrap Exploris 480 (Thermo) mass spectrometer. The peptides were separated on a 75 µm × 15 cm, 3 µm Acclaim PepMap reverse phase column (Thermo) at 300 nL/min using an UltiMate 3000 RSLCnano HPLC (Thermo) and eluted directly into the mass spectrometer. For analysis in the Fusion, parent full-scan mass spectra collected in the Orbitrap mass analyzer set to acquire data at 120,000 FWHM resolution and HCD fragment ions detected in the ion trap. For analysis in the Exploris 480, parent full-scan mass spectra acquired at 120,000 FWHM resolution and product ion spectra at 15,000 resolution. Proteome Discoverer 3.0 (Thermo) was used to search the data using SequestHT. For analysis a custom database with the ZC3H7A_440-971_ sequence was used with the human database from Uniprot. The search was limited to tryptic peptides, with maximally two missed cleavages allowed. Cysteine carbamidomethylation, methionine oxidation, and cysteine and lysine acetylation were set as variable modifications. The precursor mass tolerance was 10 ppm, and the fragment mass tolerance was 0.6 Da for data obtained on the Fusion and 0.02 Da for data obtained on the Exploris 480. The Percolator node was used to score and rank peptide matches using a 1% false discovery rate. Label-free quantitation of extracted ion chromatograms from MS1 spectra was performed using the Minora node in Proteome Discoverer.

### In vitro reaction of SAMT-247 with MGMT

His-tagged recombinant MGMT protein (Abcam, 20 µM) was incubated with either 100 µM SAMT-247 or DMSO vehicle control in THP-1 cell lysate at 37 °C for 1 h. Following incubation, MGMT was enriched by pulldown with cobalt affinity beads (Thermo Fisher Scientific). After the final wash, beads were resuspended in 1× lysis buffer containing 5% SDS in 50 mM TEAB and processed for S-Trap digestion (Protifi). Proteins were alkylated with iodoacetamide without prior reduction to preserve modifications on cysteine and digested with trypsin (Thermo Scientific) or chymotrypsin (Promega) at an enzyme to peptide ratio of 1:50 at room temperature overnight. Mass spectrometry analysis was performed using an Orbitrap Exploris 480 (Thermo) mass spectrometer. The peptides were separated on a 75 µm × 15 cm, 3 µm Acclaim PepMap reverse phase column (Thermo) at 300 nL/min using an UltiMate 3000 RSLCnano HPLC (Thermo) and eluted directly into the mass spectrometer. Parent full-scan mass spectra acquired at 120,000 FWHM resolution and product ion spectra at 15,000 resolution. Proteome Discoverer 3.0 (Thermo) was used to search the data using SequestHT with the human database from Uniprot. The search was limited to chymotryptic or tryptic peptides, with maximally two missed cleavages allowed. Cysteine carbamidomethylation, methionine oxidation, and cysteine or lysine acetylation were set as variable modifications. The precursor mass tolerance was 10 ppm, and the fragment mass tolerance was 0.02 Da. The Percolator node was used to score and rank peptide matches using a 1% false discovery rate. Label-free quantitation of extracted ion chromatograms from MS1 spectra was performed using the Minora node in Proteome Discoverer.

## Results

### Effect of SAMT-247 on unstimulated THP-1 monocyte cells

SAMT-247 is a small molecule thioester that targets the zinc-binding domains of HIV-1 NC. Recent studies on the effectiveness of SAMT-247 in rhesus macaque models of SIV infection have suggested that SAMT-247 may impact other protein targets resulting in the modulation of immune functions.^9^ As a first step to investigate new targets of SAMT-247, THP-1 monocyte cells were treated with SAMT-247 and global proteomics used to compare the effects on protein level relative to control cells treated with DMSO (Figure 2A). Among the > 4000 proteins identified and quantified, 105 had a significant (p < 0.05) change in protein level upon SAMT-247 treatment. Within this group, only six proteins showed at least a 1.15-fold change in level (Figure 2B), suggesting that treatment with SAMT-247 has minimal effects on a large part of the proteome of THP-1 cells. Within the group of six proteins that show some change, two were known to bind zinc: FUS, a DNA/RNA binding protein with a zinc finger domain, and ELAC2, a zinc phosphodiesterase. Overall, the minimal effects of SAMT-247 treatment on the protein levels are consistent with the generally low cytotoxicity of this compound in cellular and animal models.

**Figure 2.**
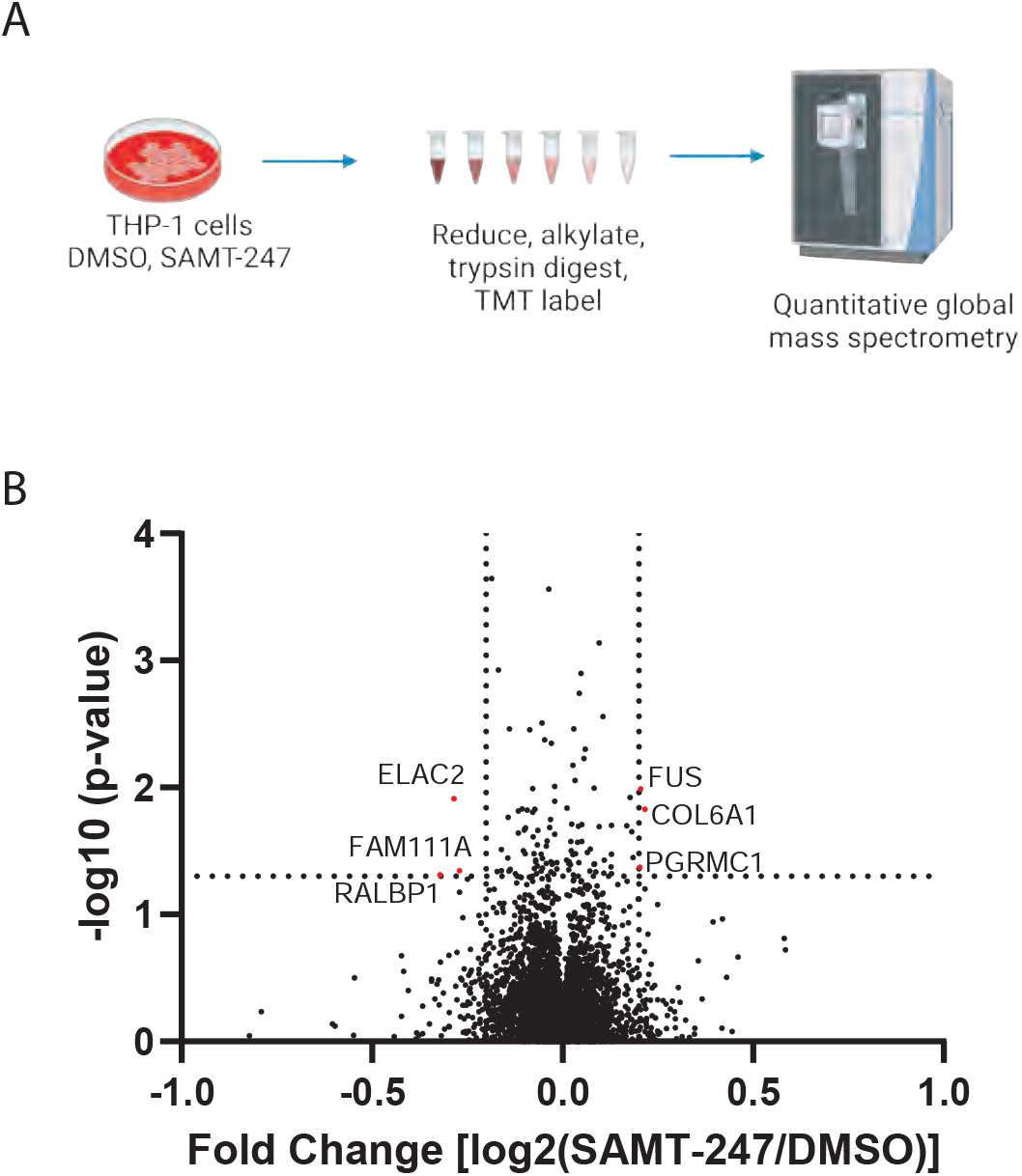
Global proteomic analysis of SAMT-247 effects. A) Schematic of global proteome analysis of effects of SMT-247 in THP-1 cells. Cells were treated with SAMT-247 or vehicle control before mass spectrometry analysis. Figure created with BioRender. B) Volcano plot of the changes in protein abundance identified in SAMT-247-treated THP-1 cells as compared with DMSO-treated cells.

Within cells, SAMT-247 promotes the transfer of an acetyl group to the sidechains of cysteine and lysine in proteins (Figure 1), leading to unfolding of the protein and aggregation.^5, 6^ To examine whether SAMT-247 destabilizes proteins other than NC, we employed a global proteomics approach to compare protein thermal stability. Thermal proteome profiling (TPP), also termed CETSA-MS, is an unbiased mass spectrometry approach that measures shifts in the thermodynamic stability of proteins by measuring protein melting points on a global scale.^18, 19^ Thus, changes in protein stability due to small molecule binding events or other types of reaction can be detected by this approach. In the context of SAMT-247, we wanted to determine whether the additional acetylation of proteins would lead to changes in protein stability across the proteome. In the experiment, cells were treated with either SAMT-247 or DMSO control, then aliquots heated to a set temperature between 37 – 67 °C. After removal of insoluble material, the soluble protein level was quantitated by isobaric tandem mass tags (TMT) (Figure 3A). From the quantitation of soluble protein at each temperature, the melting profile can be reconstructed in the presence and absence of the inhibitor. Changes in thermal stability then indicate effects of the drug.

**Figure 3.**
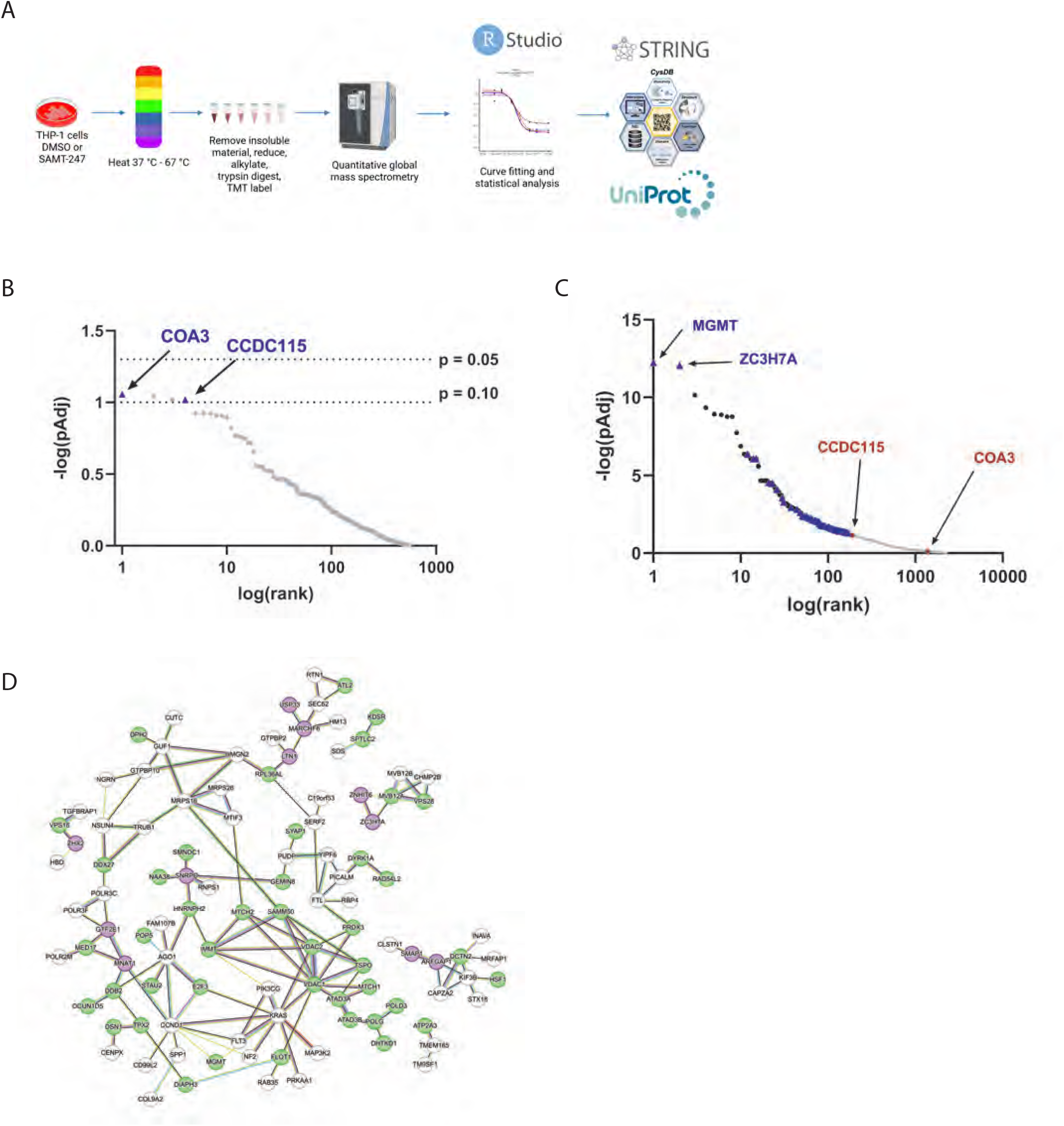
Thermal proteome profiling analysis of SAMT-247 effects in unstimulated and PMA/Iono-stimulated cells. A) Schematic of thermal proteome profiling analysis of effects of SAMT-247 in THP-1 cells. Cells were cultured, either unstimulated or stimulated with PMA/Iono, then treated with DMSO or SAMT-247. Post harvest, cells underwent lysis, differential heating, digestion, and TMT labeling, prior to quantitative mass spectrometry. Figure created with BioRender. B) Log plot illustrating Benjamini-Hochberg corrected p-value for each protein within the non-PMA/Iono-stimulated samples. Results show no protein produced a significant FDR-corrected melting shift below 0.05. Protein COA3 presents with a p value of 0.0878, and CCDC115 with 0.0956. C) Log plot illustrating Benjamini-Hochberg corrected p-value for each protein within the PMA/Iono-stimulated samples. Blue points indicate proteins denoted as containing a reactive cystine. Red points indicate two protein melt curves determined to be significantly shifted in the non-PMA/Iono stimulated cohort. Within the stimulated cohort, the significance of the difference in thermal melting for CCDC115 was p_Adj_ = 0.0726 and COA3 was p_Adj_ = 0.818. D) STRING analysis of selected significantly shifted proteins in the presence of SAMT-247 and PMA/Iono stimulation. Nodes in purple indicate proteins with zinc-finger domains, while nodes in green indicate demonstrated cysteine druggability.

TPP was first applied to THP-1 cells, as performed for the global proteome analysis. Overall, more than 4500 proteins were quantified across two biological replicates, with more than 3000 quantified in both replicates. Among these proteins, none showed a significant (p_adj_ < 0.05) change in thermal melting when treated with SAMT-247. Only a small number of proteins trended to altered stabilization with p_adj_ < 0.075 (Figure 3B). These results are consistent with the global proteome analysis that showed very few proteins were affected by SAMT-247 in THP-1 cells.

### SAMT-247 has widespread effects in the presence of immune stimulation

Our previous experiments suggested that SAMT-247 may have greater effects in stimulated immune cells. For example, cells isolated from human blood treated *in vitro* with phorbol 12-myristate 13-acetate/Ionomycin (hereafter referred to as PMA/Iono) and SAMT-247 exhibited a significantly greater trend in per-cell zinc staining intensity than cells stimulated with PMA alone, suggesting that SAMT-247 may enhance zinc availability within the system.^9^ To mimic this immune-stimulated condition, THP-1 cells were treated with PMA/Iono, causing them to differentiate into macrophages. At this stage, the cells were either treated with SAMT-247 or DMSO (control) and the TPP repeated. Analysis of the TPP data indicated a large amount of biological variation, likely due to variation in the response to PMA/Iono and the complexity of the differentiation process. Across the biological replicates, 7560 proteins were able to be identified. Of these, 5963 proteins were quantified in at least one replicate and 5210 proteins quantified in both replicates. Analysis of the proteins quantified in both biological replicates found that 170 had a significant (p_adj_ < 0.05) shift in their thermal melting profile after SAMT-247 treatment (Figure 3C). Thus, SAMT-247 had a more profound effect in the stimulated macrophages than in the unstimulated monocytes.

Of the 170 proteins that were significantly affected by SAMT-247 treatment, seven were annotated as containing a zinc-coordination domain, 64 were annotated in CysDB as having a reactive cystine, and 10 contained both.^16^ For example, the U1 small nuclear ribonucleoprotein C (U1-C), encoded by the *SNRPC* gene, is a component of the U1 spliceosomal small nuclear ribonucleoparticle snRNP (U1snRNP) that has a matrin-type zinc finger domain in its N-terminus. This protein was found to be reproducibly destabilized in PMA/Iono-stimulated THP-1 cells upon treatment with SAMT-247 (Supplemental Figure 1A). Of the remaining 89 proteins affected by SAMT-247 treatment, STRING analysis indicated that 39 are in a direct network with a zinc-coordinating protein or a protein with a reactive cysteine (Figure 3D). Therefore, these 39 proteins could be affected indirectly via an effect on an interaction parter. For example, RNPS1, a protein that does not have a zinc-binding domain or an accessible cysteine residue, was found to be significantly destabilized in the presence of SAMT-247 (Supplemental Figure 1B). This protein has been shown to interact with the U1snRNP and U1-C.^20, 21^ Comparison of the thermal melt curves of U1-C and RNPS1 shows that they are statistically similar, with analysis of all significantly shifted proteins indicating very similar deviation patterns and relative placement among the results for these two proteins (Supplemental Figure 1C). It is therefore likely that destabilization of U1-C induced by SAMT-247 subsequently destabilizes RNPS1 within the ribonucleoparticle where both proteins interact.

Among the proteins affected by SAMT-247 in the macrophages, we identified fifteen with roles in cellular metabolism. Another five proteins are important in mitochondrial organization and five more are involved in mitochondrial DNA replication and translation. For example, LYRM2, SAMM50, and NUBPL all function in assembly of respiratory chain complex I. To better understand the effect of SAMT-247 on cellular metabolism, Seahorse assays were used to examine the effects of PMA/Iono alone or PMA/Iono+SAMT-247 on cellular energy production by both mitochondrial respiration and glycolysis. Looking first at unstimulated cells, we found that there was no significant effect of SAMT-247 on mitochondrial respiration (OXPHOS), consistent with the global proteomic and TPP results (Figure 4A-G). Stimulation with PMA/Iono alone significantly decreased OXPHOS, including basal respiration, ATP production, maximal respiration, and proton leak. When used in combination, SAMT-247 amplified the effect of PMA/Iono on OXPHOS, further reducing all factors but proton leak. Treatment with SAMT-247 alone only slightly increased the glycolytic function of the cells, with small but significant increases in glycolytic capacity and glycolytic reserve of the cells (Figure 4H-L). These parameters were all increased upon stimulation with PMA/Iono alone and further enhanced when PMA/Iono-stimulated cells were treated with SAMT-247. These data indicate that the response to PMA/Iono in THP-1 cells is anti-metabolic and pro-glycolytic, consistent with M1 macrophage metabolism^22-24^, and that treatment with SAMT-247 augments these metabolic changes.

**Figure 4.**
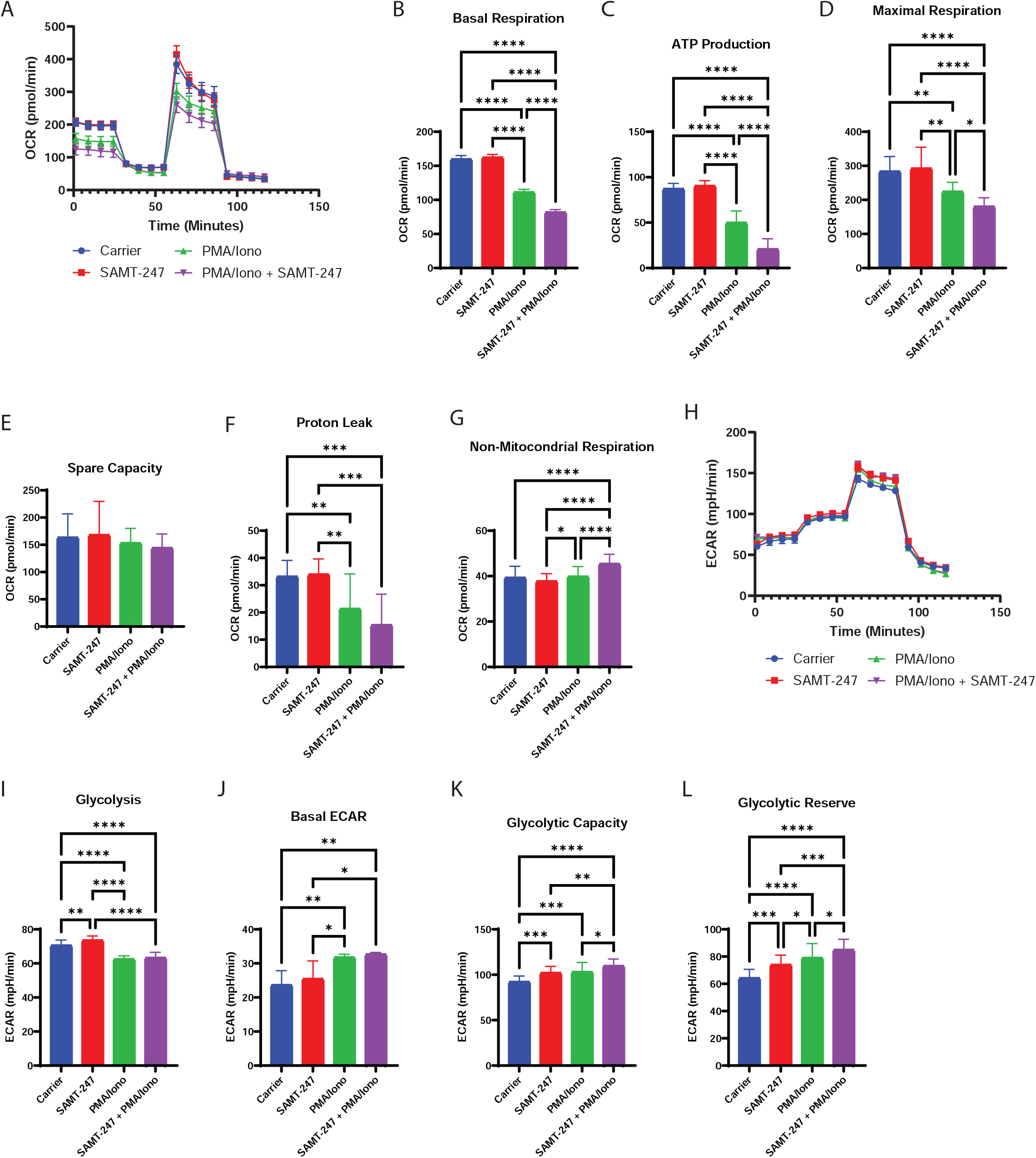
Effects of SAMT-247 on THP-1 cell metabolism. A) Mitochondrial function was measured using the Seahorse XF^e^96 Cell Mito Stress Test Kit following treatment of THP-1 cells with PMA/Iono, SAMT-247, or PMA/Iono+SAMT-247. B-G) Bioenergetic metrics were calculated from the Seahorse XF^e^96 extracellular flux bioanalyzer OCR traces shown in panel A as described in the methods. H) Glycolytic function was measured using the Seahorse XFe96 Glycolytic Stress Test Kit following treatment of THP-1 cells with PMA/Iono, SAMT-247, or PMA/Iono+SAMT-247. I-L) Bioenergetic metrics were calculated from the Seahorse XF^e^96 extracellular flux bioanalyzer ECAR traces shown in panel H as described in the methods.

### Validation of ZC3H7A and MGMT as direct reaction targets of SAMT-247

The protein most significantly affected by SAMT-247 in stimulated THP-1 cells was ZC3H7A. This zinc finger protein was reproducibly destabilized by SAMT-247 treatment in the PMA/Iono-stimulated cells (Figure 5A). ZC3H7A is comprised of four zinc finger (ZF) domains in the C-terminus (Figure 5B) and is known to bind to miR7-1, miR16-2, and miR29A hairpins, recognizing the 3′-ATA(A/T)-5′ motif in the apical loop to regulate processing. The large effect of SAMT-247 on ZC3H7A thermal stability raised the question of whether one or more of the zinc fingers could be targeted by SAMT-247.

**Figure 5.**
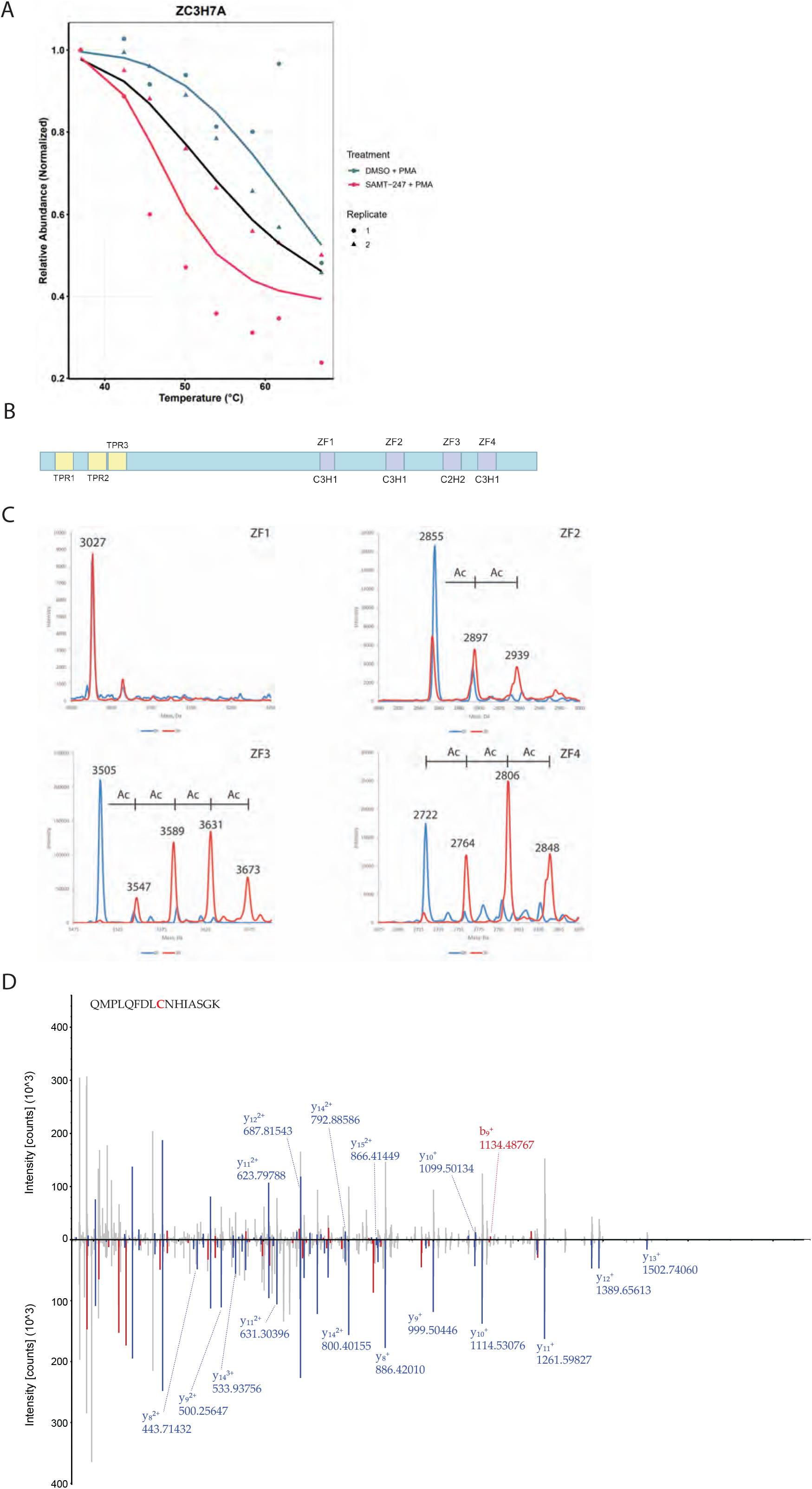
SAMT-247 reacts with the zinc fingers of ZC3H7A. A) Melting curve from TPP analysis of ZC3H7A in the absence (blue) and presence (red) of SAMT-247. Data for replicate 1 is shown as a circle and for replicate 2 as a triangle. The black line indicates the null hypothesis of no change. B) Schematic of ZC3H7A domain structure, showing the four zinc finger domains in the C-terminus. C) Reconstructed mass spectra showing reaction of ZC3H7A zinc finger peptides incubated with SAMT-247 or vehicle control. Blue spectra are DMSO-treated controls and orange spectra are treated with SAMT-247. Masses consistent with SAMT-247 acetylation are indicated. D) Butterfly plot representing the MS/MS spectra of ZC3H7A(767-782) with carbamidomethylated (bottom) or SAMT-247-modified (top) Cys775. B-(red) or y-(blue) ions indicating the mass shift are highlighted.

To first investigate whether ZC3H7A could be directly targeted by SAMT-247, mass spectrometry was used to analyze covalent modification of zinc-refolded peptides representing each of the four zinc fingers. Reaction with SAMT-247 adds a mass of 42 Da, allowing the number of reaction sites to be directly monitored. Following incubation of each peptide with a molar excess of SAMT-247, we characterized the potential reaction on each peptide. Although ZF1 did not react with SAMT-247, as demonstrated by identical spectra for the untreated and treated peptide forms, the other three peptides did show varying reaction with SAMT-247 (Figure 5C). ZF2 had two sites of covalent modification; ZF4 showed three sites of reaction; and ZF3 showed four sites.

Since isolated peptides do not always represent reactivity in the context of a full protein, the reaction of ZC3H7A_440-971_ with SAMT-247 was next analyzed. In this recombinant protein construct, all four zinc fingers of ZC3H7A are present, in addition to a large portion of the middle of the protein; only the N-terminus was removed to eliminate disordered regions. Reaction of a 5-fold molar excess of SAMT-247 with ZC3H7A_440-971_ was monitored by mass spectrometry. Following tryptic digestion to map sites of reaction, modification of multiple cysteine and lysine residues throughout the protein was observed (Figure 5D, Supplemental Figure 2). As observed for the isolated peptides, no modification within ZF1 was detected whereas modification of cysteine and lysine residues in ZF2, ZF3, and ZF4 was observed. For example, Cys775 and Cys790 in ZF2 were uniquely modified in the SAMT-247-treated samples. Acetylated forms of Cys859 in ZF3 and Cys907 and Cys915 in ZF4 were detected in both the untreated and treated samples, though unique lysine modifications in these domains in following SAMT-247-treatment suggest that further reaction in these domains occurs beyond anything that arises during purification. Multiple lysine residues outside of the zinc-coordinating domains were also found in the modified state, similar to what has previously been observed for HIV NC.^7^

Finally, reaction of ZC3H7A_440-971_ was performed in the presence of THP-1 lysate to examine whether the presence of other proteins would affect SAMT-247 reaction. Similar results were obtained where multiple sites of ZC3H7A_440-971_ modification were observed in the protein, including acetylation at a cysteine residue in ZF1 and lysine residues in ZF2 and ZF3 (Supplemental Figure 3). As shown with ZC3H7A_440-971_ alone, acetylation was also observed on lysine residues outside of the zinc finger domains. Combined, these results indicate that ZC3H7A is a bona fide target of SAMT-247.

A second highly significant protein from the TPP analysis was MGMT, a DNA damage repair protein in which a zinc ion stabilizes a helical bridge between a two-domain alpha/beta structure.^25^ Like ZC3H7A, MGMT was significantly destabilized by SAMT-247 treatment (Figure 6A). The DNA repair process catalyzed by MGMT involves a suicide reaction with an exposed cysteine residue, which has previously been shown to be accessible for reaction.^16^ Thus, we hypothesized that this residue could also be targeted by SAMT-247. Reaction of recombinant MGMT with SAMT-247 resulted in specific modification of Lys107 and increased modification of Lys165 as compared with the unreacted protein (Supplemental Figure 4). When MGMT was incubated with SAMT-247 in the presence of cellular extract, specific reaction was observed at Lys104 and Lys107 (Figure 6B,C). Thus, MGMT is also a true target of SAMT-247.

**Figure 6.**
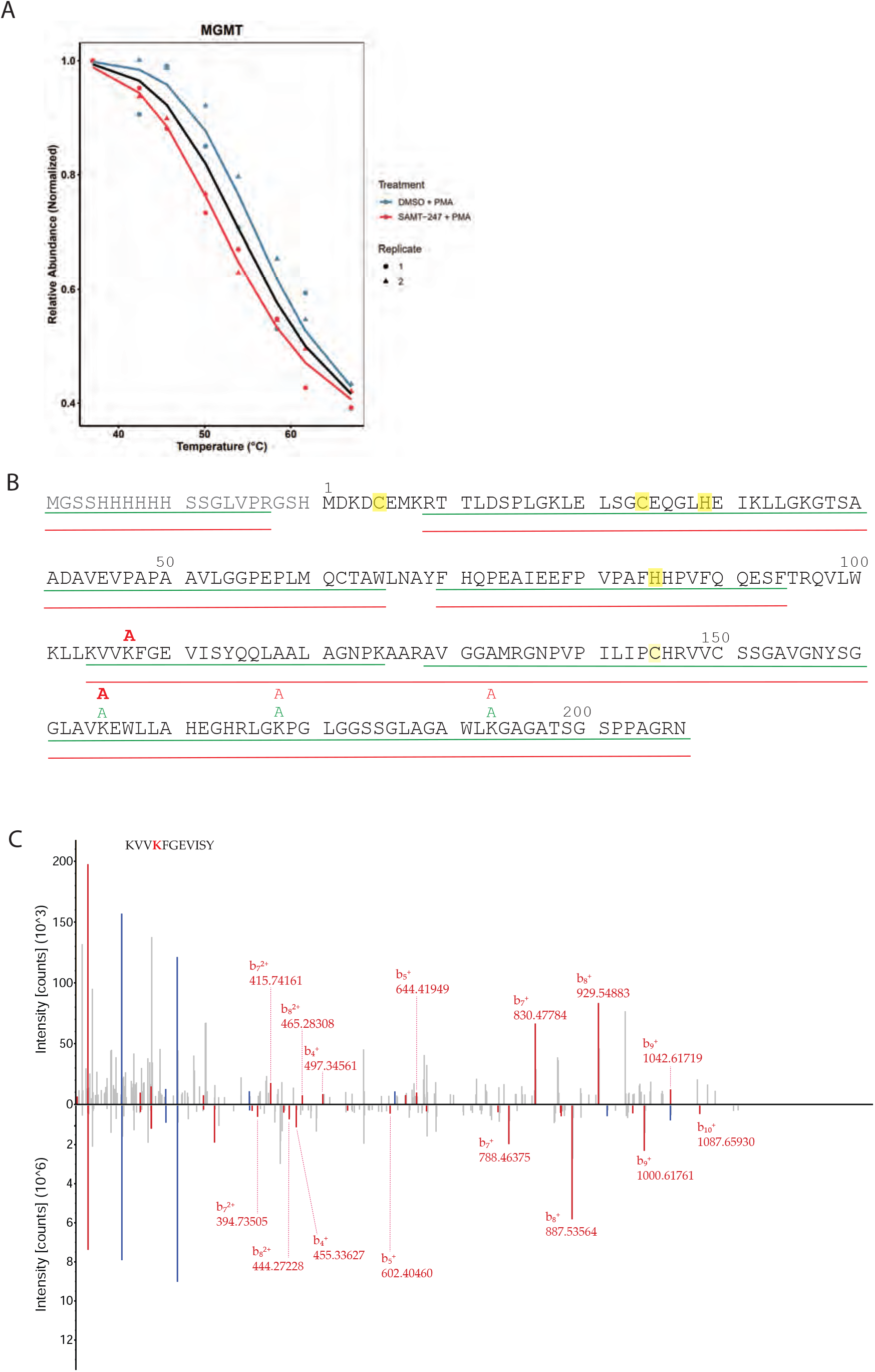
MGMT is modified by SAMT-247. A) Melting curve from TPP analysis of MGMT in the absence (blue) and presence (red) of SAMT-247. Data for replicate 1 is shown as a circle and for replicate 2 as a triangle. The black line indicates the null hypothesis of no change. B) Modification of His-MGMT in the absence or presence of SAMT-247. The green line indicates the sequence coverage of MGMT in the DMSO-treated control and the red line in the SAMT-247 reactions. Green “A” indicates identification a site of acetylation in the DMSO control and red “A” a site in the SAMT-247 reaction. Bold font represents sites with >5-fold increase in acetylation in the SAMT-247 reaction as compared to the DMSO control. C) Butterfly plot representing the MS/MS spectra of MGMT (104-114) with unmodified (bottom) or SAMT-247-modified (top) Lys107. B-ions indicating the mass shift are highlighted in red.

## Discussion

We have previously shown that SAMT-247 inhibits the HIV NC protein by covalent modification of the zinc-coordinating cysteine residues. As covalent modification is a permanent mechanism of protein inhibition, understanding the off-target effects of these inhibitors is critical. While SAMT-247 has been shown to exhibit low cytotoxicity in cellular and animal models^26-28^, this empirical observation does not preclude reactions with alternate targets. In addition to identifying potential cytotoxic targets, understanding of protein targets of SAMT-247 could promote novel applications for treatment of other diseases.

Here, we have used global proteomics approaches to investigate the cellular effects of SAMT-247 treatment and to characterize alternative targets. In unstimulated THP-1 cells, treatment with SAMT-247 showed minimal effects on global protein level and thermal stability. These results were somewhat surprising, as the structure of SAMT-247 suggests that it would be highly reactive towards free thiols. The minimal effect observed with SAMT-247 in basal THP-1 cells combined with its low cytotoxicity suggest that any SAMT-247 reactions that do occur must be quickly cleared or occur at a low enough level such that no functional consequences are observed.

In contrast, thermal proteome profiling performed in THP-1 cells stimulated with PMA/Iono identified 170 affected proteins. Within the group of affected proteins, nearly half have annotated zinc fingers or have previously been shown to have reactive cysteine residues.^16^ The mechanism of action of SAMT-247 targeting HIV NC begins with covalent modification of Cys36. Although this is a zinc-coordinating residue, it has been shown to exist in an equilibrium with a short-lived state where it does not coordinate zinc; it is this state that is thought to be the target of SAMT-247. Once the initial reaction and acetylation of the cysteine sidechain occurs, zinc coordination within the finger is destabilized, facilitating reaction at other sites in the domain. Since SAMT-247 is known to react with other zinc-coordination motifs to varying degrees,^12^ the observation of 17 proteins with zinc finger domains among the reaction targets was an expected finding. Likewise, proteins with accessible cysteine residues are reasonable candidates for reaction. It is interesting to compare the overall low effects of SAMT-247 between unstimulated and PMA/Iono-stimulated cells.

As demonstrated here, the PMA/Iono-stimulated THP-1 cells have higher glycolytic activity (Figure 4); this result is consistent with previous results for activated immune cells.^29^ High glycolytic activity is correlated with increased acetyl-CoA levels, as it is produced by glycolysis and is a precursor to the citric acid cycle.^30^ These increased acetyl-CoA levels may result in better SAMT-247 regeneration via recycling reaction with acetyl-CoA, thereby leading to greater activity of the inhibitor. Additionally, as histone acetylation is increased in macrophages as compared with monocytes^31^, PMA/Iono-induced differentiation of the THP-1 cells may lead to increased transcriptional activity. Thus, greater SAMT-247 regeneration in transcriptionally-active cells may result in the ability of the inhibitor to target more proteins, particularly those with zinc-binding domains.

PMA/Iono functions primarily by activating protein kinase C, leading to upregulation of cytokines and other myeloid cell activation effects. Our results suggest that SAMT-247 may have an unexplored efficacy against inflammation or autoimmune disorders. In an earlier study, we showed that SAMT-247 treatment was associated with a significant decrease in TNF-α production in Th1 and Th2 cells stimulated with PMA/Iono, as well as increased IL-10 production in PMA/Iono-stimulated Th1 cells.^9^ Zinc homeostasis is known to be critical for immune regulation^32^ and several zinc finger proteins have been shown to play important roles in the immune response.^33^ Thus, modulation of zinc fingers and zinc homeostasis by SAMT-247 could have important implications for immune signaling.

Among the proteins affected by SAMT-247 in the PMA/Iono-stimulated THP-1 cells, we demonstrated that two are targeted by the inhibitor *in vitro*. ZC3H7A is comprised of four zinc-finger domains, three with C_3_H_1_ coordination and one with C_2_H_2_ coordination. Using isolated peptides and whole protein, we found that residues in the first three zinc finger domains could be modified by SAMT-247. Like HIV NC, ZC3H7A is an RNA-binding protein; it regulates microRNA biogenesis, and binds to apical loops in the hairpins of miR-7-1, miR-16-2, and miR-29a.^34^ It has been shown to preferentially bind and destabilize mRNAs enriched in non-optimal codons.^35^ Similar to the reaction of SAMT-247 with NC, it is likely that SAMT-247 acetylation within the zinc-finger domains of ZC3H7A would destabilize their fold and thereby inhibit RNA binding. Consistent with the importance of the zinc fingers for overall protein fold, SAMT-247 treatment resulted in destabilization of ZC3H7A by thermal melting. Interestingly, ZC3H7A was identified in whole-blood transcriptomics as the only gene negatively associated with the efficacy of an HIV vaccine candidate tested in young non-human primates.^36^ The ability of SAMT-247 to destabilize ZC3H7A protein may help explain its synergistic effect in a similar vaccine strategy observed in a subsequent non-human primate vaccine study.^9^

MGMT is a DNA repair protein that repairs *O*^6^-alkylguanoine by a stoichiometric transfer of the alkyl group to an active site cysteine, thereby irreversibly inactivating the protein and targeting it for degradation.^37^ As such, it also promotes tumor resistance to alkylating agents used as chemotherapeutics for several cancer types, including glioblastoma and malignant melanoma.^38^ Combination of MGMT inhibitors with alkylating chemotherapeutics increases the effectiveness in tumor cells, but leads to toxicity in normal tissues.^39^ Given the low activity of SAMT-247 in non-stimulated THP-1 cells observed here, it is possible that its use could represent a new means to prevent MGMT resistance with lower toxicity in healthy cells. Indeed, we have previously shown that SAMT-247 combines synergistically with both vaccines and antiretrovirals for inhibition of HIV infection with no effect on toxicity, suggesting that it could be combined with alkylating chemotherapeutics to increase their efficacy as well.^8, 9, 11^ Further, as the thiol resulting from SAMT-247 reaction is intracellularly re-acylated to form a new active thioester, each reaction with MGMT would inactivate a single molecule of the protein but not necessarily of the drug, potentially allowing it to have greater overall potency and/or be used at a lower concentration.

Here, we have used proteomics approaches to investigate the overall reactivity of SAMT-247, demonstrating that it has minimal effects in unstimulated THP-1 cells but has multiple potential reaction targets once those cells were stimulated with PMA/Iono. Many of those proteins contain zinc-binding domains or reactive cysteine residues, representing likely bona fide targets. The difference in effect in unstimulated and stimulated cells is likely due to differential protein expression or stability in the two cellular conditions, suggesting a potential therapeutic window for SAMT-247 utilization. The results of this study further highlight the potential of this inhibitor to be repurposed as a treatment for other diseases, including cancer, either alone or in combination with other therapeutics. For example, the confirmed interaction between SAMT-247 and MGMT could be useful for future chemotherapeutic use. MGMT-mediated resistance to temozolomide is a significant roadblock to effective chemotherapeutic treatment in glioblastoma multiforme.^40^ Post-translational targeting and inactivation of MGMT by SAMT-247 may hinder the de-alkylation of cellular DNA, improving apoptotic efficiency. Further characterization of these new reaction targets and the effects of SAMT-247 modification *in vivo* is a critical next step in the development of this molecule.

## Acknowledgements

This research was supported by the Intramural Research Program of the Center for Cancer Research, National Cancer Institute and National Institute of Diabetes and Digestive and Kidney Diseases, National Institutes of Health. The contributions of the NIH author(s) are considered Works of the United States Government. The findings and conclusions presented in this paper are those of the author(s) and do not necessarily reflect the views of the NIH or the U.S. Department of Health and Human Services.

## Data Availability

The mass spectrometry proteomics data have been deposited to the ProteomeXchange Consortium via the massIVE partner repository with the dataset identifier PXD077667.

## Notes

### Competing Interest Statement

The authors have declared no competing interest.

