## Supplemental Figure 4 for "Global Proteomics Investigation of SAMT-247 Targets: An Antiviral Thioester that Acetylates Zinc Finger Proteins"

**Supplemental Figure 4.** Modification of His-MGMT in the absence or presence of SAMT-247. The green line indicates the sequence coverage of His-MGMT in the DMSO-treated control and the red line in the SAMT-247 reactions. Green "A" indicates identification a site of acetylation in the DMSO control and red "A" a site in the SAMT-247 reaction. Bold font represents sites with >5-fold increase in acetylation in the SAMT-247 reaction as compared to the DMSO control.

Supplemental Figure 4

```
                     1                                                      50
MGSSHHHHHH SSGLVPRGSH MDKDCEMKRT TLDSPLGKLE LSGCEQGLHE IKLLGKGTSA ADAVEVPAPA AVLGGPEPLM QCTAWLNAYF HQPEAIEEFP VPAFHHPVFQ

                                                                                 A             A                A
        100      A                                                  150          A             A                A      200
QESFTRQVLW KLLKVVKFGE VISYQQLAAL AGNPKAARAV GGAMRGNPVP ILIPCHRVVC SSGAVGNYSG GLAVKEWLLA HEGHRLGKPG LGGSSGLAGA WLKGAGATSG SPPAGRN
```

untreated

+ 5-fold SAMT-247
