## Supplemental Figure 3 for "Global Proteomics Investigation of SAMT-247 Targets: An Antiviral Thioester that Acetylates Zinc Finger Proteins"

**Supplemental Figure 3.** Modification of ZC3H7A$_{440\text{-}971}$ incubated in THP-1 lysate in the absence or presence of SAMT-247. The red line indicates the sequence coverage of ZC3H7A$_{440\text{-}971}$ in the SAMT-247 reaction. Red “A” indicates sites of acetylation, consistent with SAMT-247 reaction. Yellow highlighting indicates the four zinc finger domains.

Supplemental Figure 3

```
                             A                 A
HELRQACQIC FVKSGPKLMD FTYHANIDHK CKKDILIGRI KNVEDKSWKK

                              A
IRPRPTKTNY EGPYYICKDV AAEEECRYSG HCTFAYCQEE IDVWTLERKG

                         A                   A    A
AFSREAFFGG NGKINLTVFK LLQEHLGEFI FLCEKCFDHK PRMISKRNKD

NSTACSHPVT KHEFEDNKCL VHILRETTVK YSKIRSFHGQ CQLDLCRHEV

   A                                          A
RYGCLREDEC FYAHSLVELK VWIMQNETGI SHDAIAQESK RYWQNLEANV

                              A                    A
PGAQVLGNQI MPGFLNMKIK FVCAQCLRNG QVIEPDKNRK YCSAKARHSW

                            A           A                A
TKDRRAMRVM SIERKKWMNI RPLPTKKQMP LQFDLCNHIA SGKKCQYVGN

CSFAHSPEER EVWTYMKENG IQDMEQFYEL WLKSQKNEKS EDIASQSNKE

                                           A              A
NGKQIHMPTD YAEVTVDFHC WMCGKNCNSE KQWQGHISSE KHKEKVFHTE

DDQYCWQHRF PTGYFSICDR YMNGTCPEGN SCKFAHGNAE LHEWEERRDA

           A              A
LKMKLNKARK DHLIGPNDND FGKYSFLFKD LN
```
