## Supplemental Figure 2 for "Global Proteomics Investigation of SAMT-247 Targets: An Antiviral Thioester that Acetylates Zinc Finger Proteins"

**Supplemental Figure 2.** Modification of ZC3H7A$_{440-971}$ in the absence or presence of SAMT-247. The green line indicates the sequence coverage of ZC3H7A$_{440-971}$ in the DMSO-treated control and the red line in the SAMT-247 reactions. Green "A" indicates identification a site of acetylation in the DMSO control and red "A" a site in the SAMT-247 reaction. Bold font represents sites with >5-fold increase in acetylation in the SAMT-247 reaction as compared to the DMSO control. Yellow highlighting indicates the four zinc finger domains.

Supplemental Figure 2

```
             A                 A   A        A    A
HELRQACQIC FVKSGPKLMD FTYHANIDHK CKKDILIGRI KNVEDKSWKK

                  A                                 A
IRPRPTKTNY EGPYYICKDV AAEEECRYSG HCTFAYCQEE IDVWTLERKG

                                     A
AFSREAFFGG NGKINLTVFK LLQEHLGEFI FLCEKCFDHK PRMISKRNKD

           A
NSTACSHPVT KHEFEDNKCL VHILRETTVK YSKIRSFHGQ CQLDLCRHEV

                                          A
RYGCLREDEC FYAHSLVELK VWIMQNETGI SHDAIAQESK RYWQNLEANV

                  A
                  A A                  A  A
PGAQVLGNQI MPGFLNMKIK FVCAQCLRNG QVIEPDKNRK YCSAKARHSW

                            A
                A          AA         A       AA
TKDRRAMRVM SIERKKWMNI RPLPTKKQMP LQFDLCNHIA SGKKCQYVGN

                 A
A                A                                  A
CSFAHSPEER EVWTYMKENG IQDMEQFYEL WLKSQKNEKS EDIASQSNKE

  A                 A     A
  A                 A   A A      A          A
NGKQIHMPTD YAEVTVDFHC WMCGKNCNSE KQWQGHISSE KHKEKVFHTE

                  A        A
                  A        A       A
DDQYCWQHRF PTGYFSICDR YMNGTCPEGN SCKFAHGNAE LHEWEERRDA

                        A
LKMKLNKARK DHLIGPNDND FGKYSFLFKD LN
```

untreated

+ 5-fold SAMT-247
