## Supplemental Figure 1 for "Global Proteomics Investigation of SAMT-247 Targets: An Antiviral Thioester that Acetylates Zinc Finger Proteins"

**Supplemental Figure 1.** A) Melting curve from TPP analysis of RU1C in the absence (blue) and presence (red) of SAMT-247. Data for replicate 1 is shown as a circle and for replicate 2 as a triangle. The black line indicates the null hypothesis of no change. B) Melting curve from TPP analysis of RNPS1 in the absence (blue) and presence (red) of SAMT-247. Data for replicate 1 is shown as a circle and for replicate 2 as a triangle. The black line indicates the null hypothesis of no change. C) Residual sum of squares analysis of all significantly shifted proteins.

Supplemental Figure 1

A

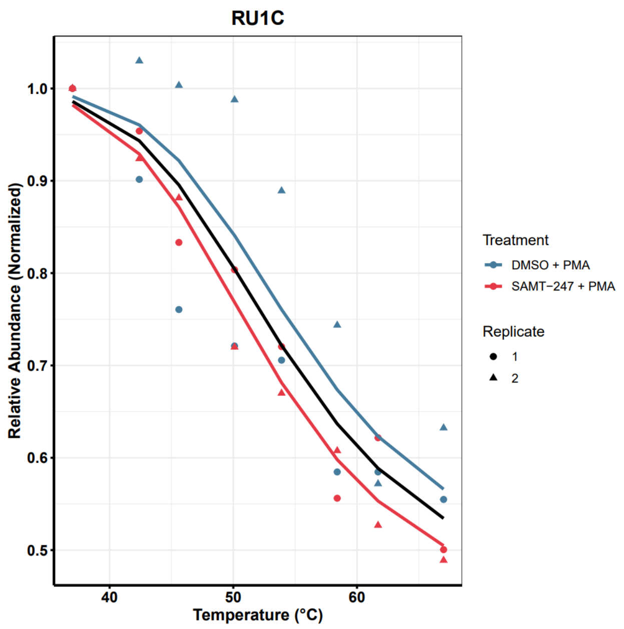


B

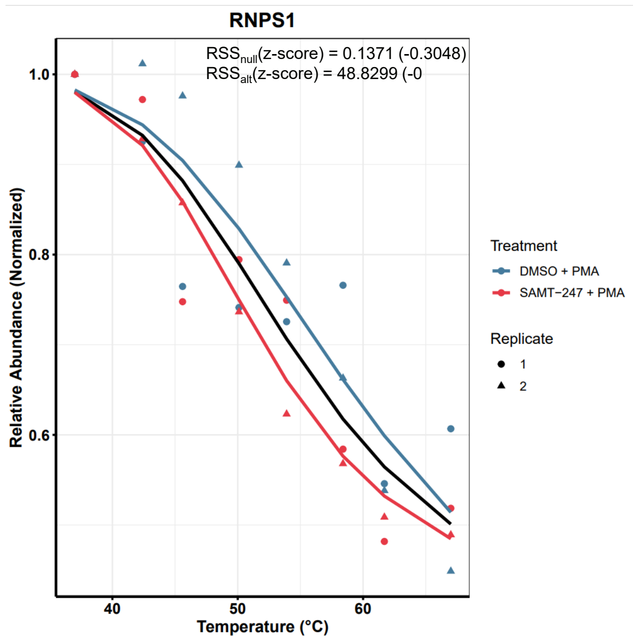


C

| Reactive | Non-reactive | $RSS_{null, z\text{-}score}$ | $\Delta RSS_{alt, z\text{-}score}$ |
|---|---|---|---|
| DDB2 | AGO1 | 0.073151 | 0.086975 |
| POP5 | AGO1 | 0.021759 | 0.040055 |
| HNRNPH2 | AGO1 | 0.062563 | 0.063618 |
| ATP2A3 | TMEM165 | 0.044688 | 0.040663 |
| DCTN2 | CAPZA2 | 0.003414 | 0.004971 |
| MVB12A | CHMP2B | 0.009969 | 0.010201 |
| DPH2 | GUF1 | 0.015809 | 0.028246 |
| FLOT1 | RAB35 | 0.028221 | 0.027701 |
| PRDX3 | FTL | 0.064912 | 0.085739 |
| GTF2E1 | POLR3C | 0.004177 | 0.009862 |
| ZHX2 | HBD | 0.008072 | 0.045685 |
| MARCHF6 | HM13 | 0.006819 | 0.015847 |
| RPL36AL | HMGN2 | 0.095891 | 0.082728 |
| MARCHF6 | SEC62 | 0.023724 | 0.023671 |
| SAMM50 | MRPS16 | 0.03459 | 0.005604 |
| MVB12A | MVB12B | 0.056408 | 0.077117 |
| VPS28 | MVB12B | 0.045948 | 0.02743 |
| SYAP1 | PUDP | 0.039878 | 0.044195 |
| SNRPC | RNPS1 | 0.037887 | 0.049313 |
| VPS16 | TGFBRAP1 | 0.036272 | 0.033474 |
| VDAC2 | TSPO | 0.076009 | 0.080874 |
